# Dynamic Adaptation in Extant Porins Facilitates Antibiotic Tolerance in Energetic *Escherichia coli*

**DOI:** 10.1101/2024.03.07.583920

**Authors:** Sayak Mukhopadhyay, Romit Bishayi, Aakansha Shaji, Annie H. Lee, Rachit Gupta, Mohammad Mohajeri, Aditya Katiyar, Brendan McKee, Isabella R. Schmitz, Rachel Shin, Tanmay P. Lele, Pushkar P. Lele

**Author notes:** **Correspondence:** Pushkar P. Lele, Artie McFerrin Department of Chemical Engineering, 241 Jack E. Brown Building, Texas A&M University, College Station, TX-77480-3122, USA. These authors contributed equally (Romit Bishayi, Aakansha Shaji).

## Abstract

Bacteria can tolerate antibiotics despite lacking the genetic components for resistance. The prevailing notion is that tolerance results from depleted cellular energy or cell dormancy. In contrast to this view, many cells in the tolerant population of Escherichia coli can exhibit motility – a phenomenon that requires cellular energy, specifically, the proton-motive force (PMF). As these motile-tolerant cells are challenging to isolate from the heterogeneous tolerant population, their survival mechanism is unknown. Here, we discovered that motile bacteria segregate themselves from the tolerant population under micro-confinement, owing to their unique ability to penetrate micron-sized channels. Single-cell measurements on the motile-tolerant population showed that the cells retained a high PMF, but they did not survive through active efflux alone. By utilizing growth assays, single-cell fluorescence studies, and chemotaxis assays, we showed that the cells survived by dynamically inhibiting the function of existing porins in the outer membrane. A drug transport model for porin-mediated intake and efflux pump-mediated expulsion suggested that energetic tolerant cells withstand antibiotics by constricting their porins. The novel porin adaptation we have uncovered is independent of gene expression changes and may involve electrostatic modifications within individual porins to prevent extracellular ligand entry.

## Introduction

Bacteria are remarkably adept at adapting to environmental stresses. Their ability to adapt to antibiotics carries significant health and economic implications (*1–8*). Even when lacking resistance genes, subpopulations of antibiotic-sensitive bacteria can exhibit phenotypic resistance, tolerating lethal doses of antibiotics for an extended period (*9–11*). Although their growth is usually arrested in the presence of the antibiotic, the tolerant cells can resume growth once the stress subsides (*12–16*). Unlike genotypic resistance, antibiotic tolerance is transient and less likely to be inherited (*17–20*). However, the presence of a tolerant pool of cells increases the likelihood of the eventual emergence of inheritable resistance (*21–23*). While genotypic resistance mechanisms are known (*24–28*), antibiotic tolerance remains obscure, partly owing to the diversity in adaptation mechanisms.

The major mechanisms of adaptation to antibiotic stress often involve decreasing the intracellular accumulation of the antibiotic. One such mechanism in Gram-negative bacteria decreases the number of porins in the outer membrane to limit the entry of antibiotics (*29–31*) (**Fig 1**). Another powerful adaptation mechanism intrinsic to many bacterial species involves efflux pumps that expel the antibiotic (**Fig 1**). These pumps utilize the proton-motive force (PMF), which also powers cellular growth and several functions, such as motility and ATP synthesis (*32, 33*).

**Fig 1.**
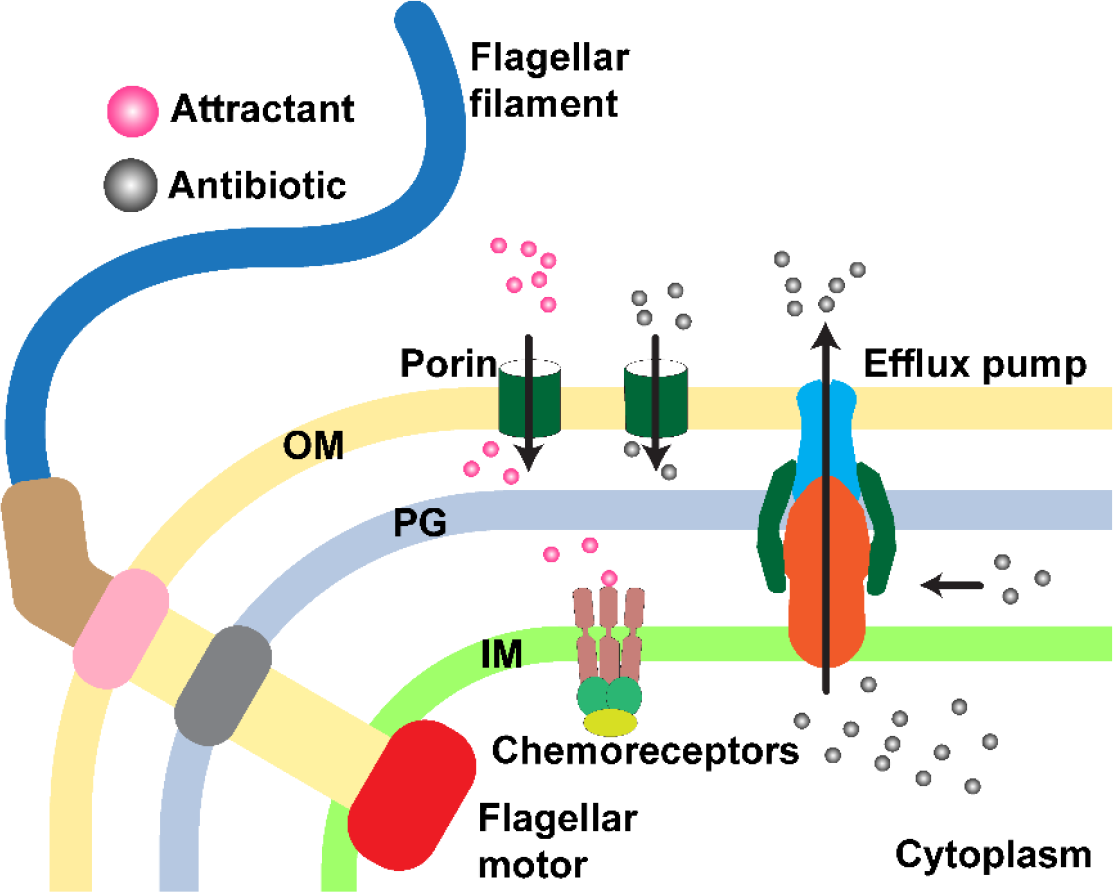
Molecular transport across gram-negative bacterial membrane. The PMF powers the rotation of the flagellar motor and active removal of antibiotics by efflux pumps. Like antibiotics, serine (attractant) diffuses through the porins in the outer membrane. In the periplasm, serine interacts with the chemoreceptors situated in the inner membrane. The chemoreceptors modulate the direction of flagellar motor rotation to mediate chemotaxis.

The rates of adaptation achievable with the porins and efflux pumps differ greatly in antibiotic-sensitive bacteria. The adaptation in porin numbers under antibiotic stress is gradual and occurs through the downregulation of porin expression. However, mere inhibition of porin synthesis does not eliminate existing porins in the cell. For this strategy to be effective, the cell must repeatedly divide during antibiotic stress to dilute the number of porins. However, this is improbable as the cell is likely to perish in the process. In contrast, efflux function can adapt within a few minutes, in principle, through the dissipation of PMF. Given the fast response afforded by efflux function, tolerance mechanisms are likelier to involve the modulation of efflux activity. Interestingly, tolerance is often associated with cellular dormancy, and dormant cells lack the PMF and ATP. Low PMF renders efflux pumps ineffective. Thus, the roles of these key processes in antibiotic tolerance remain unclear.

Brief exposures to bacteriostatic compounds that inhibit cellular respiration, which powers the PMF, can induce cell dormancy (*34*). The lack of energy stymies many processes targeted by the antibiotics, including cell wall growth. For example, a 30-minute exposure to rifampicin can induce tolerance in *Escherichia coli* to various bactericidal compounds, including ampicillin. Although rifampicin is a transcriptional inhibitor, it also dissipates cellular respiration, depriving the cell of energy. The rifampicin exposure does not merely select a stochastically occurring tolerant subpopulation; *E. coli* populations that have not been potentiated with rifampicin show negligible tolerance to bactericides (*34*).

Remarkably, our recent results demonstrate that almost 30-40% of tolerant *E. coli* cells generated by the rifampicin treatment retain motility in the presence of ampicillin and other antibiotics for several hours (*32*). This suggests that not all tolerant cells lack cellular energy, consistent with previous studies (*35–38*). It is likely that the motile-tolerant *E. coli* maintain high PMF and employ efflux pumps to survive the brief exposure to lethal doses of rifampicin, as well as the subsequent encounter with lethal concentrations of ampicillin and other antibiotics. This hypothesis is untested, and the survival mechanism of the motile-tolerant population remains a major question, underscored by motility’s crucial role in infection and fitness (*39–43*).

Probing the survival mechanisms in motile-tolerant cells is challenging, as population-based techniques such as gene expression profiling cannot distinguish between motile and nonmotile cells. To test different hypotheses, it is necessary to isolate the motile cells from the tolerant population and study them. However, conventional bulk separation methods have limited effectiveness in segregating motile and non-motile cells (*44, 45*). Single-cell assays are appropriate, but they typically involve immobilizing the cells on glass slides for microscopy. This eliminates motility, preventing discrimination between motile and non-motile cells.

Here, we employed a micro-confinement approach with a microfluidic device, known as the mother machine (*46*), to isolate motile cells from heterogeneous populations and perform single-cell assays. The device comprises a main trench flanked by orthogonal channels, with widths similar to individual bacterial cells (**Fig S1**). Typically, centrifugal forces are applied to load cells into the orthogonal channels of the device for microscopy measurements (*47*). However, this approach is unsuitable for our studies as the motile and non-motile cells cannot be distinguished once they have entered the channels. Instead, we discovered that in the absence of external forcing, only the motile cells in heterogeneous bacterial populations could overcome the heightened viscous drag and penetrate the orthogonal channels, owing to the propulsive forces they generated with their flagella. The non-motile cells were occluded from the orthogonal channels, as the confined regions prevented their diffusive entry.

We utilized micro-confinement to segregate the motile-tolerant populations generated by rifampicin treatment. Single-cell growth measurements, fluorescence microscopy, and single-cell chemotaxis assays revealed that these cells inhibited the transport of extracellular compounds through their porins within a few minutes in response to Rifampicin stress, enabling survival during subsequent exposure to Ampicillin. On the other hand, the non-motile-tolerant cells appeared to rely on cell dormancy for survival.

## Results

### Micro-channels are selectively occupied by motile cells

We cultured wild-type *E. coli* cells and resuspended them in motility buffer (MB) solutions at an optical density of 600 nm (OD_600_) of 1.3-1.5. We then introduced the cells into the microfluidic chamber. The cells were observed on a phase microscope approximately 15 min after loading the cells. The device consisted of numerous orthogonal channels measuring 25 μm in length (**Fig. S1**). Each orthogonal channel was 1.4 μm in width and 1.5 μm in height (see Supplementary text for fabrication details). The increased drag due to the channel walls hinders the entry of bacteria undergoing pure diffusion. Therefore, centrifugation has traditionally been used to force bacteria into the channels (*47*). However, we discovered that motile cells readily entered the orthogonal channels, penetrating the entire 25 μm depth without any external forcing (**Fig 2A** and **Movie 1**). We scanned several distinct regions within the device and observed cells frequently enter and exit the channels. Hence, we skipped the centrifugation step entirely in our study.

**Figure 2.** Motile cells segregate in confined channels. **A)** *Left panel*: The phase contrast image shows motile cells entering the orthogonal channels of the microfluidic device. The white arrows indicate cell trajectories over 5 seconds. *Middle panel*: Out of the total channels inspected (n = 622 channels), 68.1 ± 5.5 % were occupied by one or more motile cells of the wild-type strain. *Right panel*. The force developed by the CCW-rotating flagellar bundle helps the motile cell (yellow) easily penetrate the orthogonal channel. A switch in the direction of flagellar rotation can cause the cell to swim outward. Non-motile cells (blue) are acted upon by random Brownian forces in all possible directions. **B)** *Left panel*: Motile cells expressing eYFP penetrated the orthogonal channels of the microfluidic device. Very few cells continued swimming in the main trench. *Right panel*: Non-motile cells expressing eCFP can be observed exclusively in the main trench. **C)** Cells of the smooth-swimming mutant occupied 76.5 ± 13.9% of the channels visualized in the yellow fluorescence channel (n = 596 channels). In separate experiments, non-motile cells occupied 0% of the channels visualized in the cyan fluorescence channel (n = 537). The symbols indicate the mean occupancies by motile (black) and non-motile (white) cells in different regions observed across multiple devices (**** indicates P < 0.0001 by Welch’s t test). **D)** The composite image displays an overlay of the phase and fluorescent (yellow) channels for a 1:1 mixture of non-motile (cyan) and motile (yellow) cells. Smooth-swimming cells expressing eYFP are visible mostly in the orthogonal channels, with a few cells in the main trench. The non-motile cells expressing eCFP are exclusively visible in the main trench. In this mixture, 77.5 ± 11.9% were occupied by the motile cells, whereas 0.3 ± 0.5% of the channels were occupied by non-motile cells (n = 628 channels, **** indicates P < 0.0001 by Welch’s t test).

We employed epifluorescence microscopy to quantify how effectively the motile cells occupied the orthogonal channels. We expressed eYFP (yellow fluorescent protein) in the wild-type strain and introduced them into the device. After 15 min, we acquired fluorescence images with a sensitive sCMOS camera by suitably illuminating the cells (*Materials and Methods*). We manually counted the number of loaded and empty channels in multiple regions within the device. Even though the cells did frequently exit the channels, we observed that ∼68% of channels were occupied by wild-type cells at any given instant (**Fig 2A**, middle panel).

*E. coli* cells swim by rotating multiple flagella with the aid of ∼ 50 nm rotary motors (*48*). The motors are powered by the PMF and generate a total flagellar thrust of ∼ 0.5-1 pN on each cell (*49*). This force was adequate to penetrate the orthogonal channels in our device. Nevertheless, the flagella can switch their direction of rotation from default counterclockwise (CCW) to clockwise (CW), which can reverse the direction of propulsive force inside constricted spaces (**Fig 2A**, right panel). As a result, wild-type cells enter and escape constricted spaces readily, including the pores in soft materials such as agar (*50*). Indeed, we saw cells exit the orthogonal channels frequently, which made it challenging to monitor the properties of single cells over long durations.

Therefore, we examined whether mutant cells lacking the ability to switch the direction of rotation of their flagella were more likely to remain trapped in the channels. We employed a variant of *E. coli* RP437 deleted for the *cheY* allele (*51*). CheY, in its phosphorylated form, is a response regulator that modulates flagellar switching. In its absence, the flagella rotate exclusively CCW, and the cells swim smoothly without tumbling (*52*). We expressed eYFP in this smooth-swimming strain and loaded the cells into the microfluidic device. We appropriately illuminated the cells and imaged them in the yellow channel. In a separate set of experiments, we employed a smooth-swimming strain that expressed eCFP and was deleted for the *fliC* allele. FliC are flagellin proteins that assemble into the extracellular flagellar filaments, and in their absence, the cells are non-motile. We loaded the non-motile cells into the device and imaged them in the cyan channel.

Fluorescence imaging revealed that the smooth-swimming motile cells populated the orthogonal channels similar to the wild-type, with only a handful of cells remaining in the main trench (*Left panel*, **Fig 2B**). In contrast, cells of the non-motile strain were exclusively observed in the main trench (*Right panel*, **Fig 2B**). Remarkably, out of the several hundred channels we scanned, almost no channels were occupied by the non-motile cells (**Fig. 2C**). In comparison, ∼ 77% of the channels were occupied by the smooth-swimming cells, which was slightly higher than the corresponding number for the wildtype.

Next, we determined if the smooth-swimming cells remained trapped in individual channels for long durations. To do this, we recorded images at 5, 10, 15, 30, and 60 min after loading the cells into the device. From the images, we quantified the time-varying changes in the cell density per channel by visually tracking the number of cells in each channel. The cell density remained constant at ∼4 cells per channel (**Fig S2**). Several smooth-swimming cells were observed to enter the channels and remain trapped for the entire duration of observation. Thus, the smooth-swimming cells appear better at occupying the orthogonal channels of the microfluidic device compared to the wild-type cells.

As the orthogonal channels were stably and exclusively occupied by the motile cells (**Fig 2B**), we hypothesized that these orthogonal channels could separate a mixture of non-motile and smooth-swimming cells. We tested the hypothesis by mixing equal volumes of the two cultures. We loaded the mixture into the microfluidic device and imaged them in the dual-fluorescence channels (*Left panel*, **Fig. 2D**). The motile cells occupied ∼77.5% of the total channels, whereas the non-motile cells occupied only 2 out of 628 channels (*Right panel*, **Fig 2D**). This indicates that microchannels can effectively isolate motile cells from a mixture of motile and non-motile cells.

In conclusion, motility enables single cells to penetrate otherwise inaccessible interiors of the microfluidic device, with the depth of penetration probably limited only by the channel length. In comparison, the entry of diffusive non-motile cells into orthogonal channels appears to be kinetically limited.

### Motile-tolerant cells remain culturable

Previous studies reported that exposing cells of *E. coli* to lethal levels of rifampicin for a short time increased the likelihood of tolerance to other antibiotics (*53*). The traditional view was that rifampicin promotes tolerance by decreasing cellular respiration rates (*34*), dissipating the membrane potential (ΔΨ) —an essential component of the PMF and ATP levels. The loss of cellular energy causes metabolic dormancy, enabling the cells to withstand the effects of antibiotics, such as ampicillin, targeting active growth processes. Interestingly, the tolerant cells remain culturable and frequently resume their growth once the antibiotic levels subside (*53*).

Our recent study suggests that rifampicin induces a bifurcation of the tolerant population: nearly one out of three tolerant cells continued to swim rapidly in lethal concentrations of ampicillin for several hours, while the rest were nonmotile (*32*). As motility requires the PMF, which also produces ATP during aerobic respiration, this suggested that a significant fraction of the tolerant cells might not be dormant. We could not decisively determine in the previous work whether these motile cells remained culturable or if some of these cells belonged to a group of bacteria known as viable but non-culturable (VBNC) cells (*54, 55*). This distinction is critical in determining the mechanisms by which the motile tolerant cells survive and the relative importance of motility in bacterial persistence (*56, 57*).

To determine if the motile-tolerant cells are VBNCs, we utilized the orthogonal channels to isolate them. First, we treated the smooth-swimming strain with a lethal concentration of rifampicin for 30 min. We washed the cells in the growth media to remove the rifampicin, following earlier protocols (*32, 53*). The cells were then suspended in lethal concentrations of ampicillin for three hours (see *Materials and Methods*). A significant fraction of the cells tolerated the ampicillin exposure and several cells retained motility, consistent with earlier studies (*32, 34, 53*). We again washed the tolerant cells to remove ampicillin and loaded them into the microfluidic chamber while simultaneously observing the channels. The channels came to be exclusively occupied by the motile-tolerant cells, whereas a mix of motile and non-motile-tolerant cells remained in the main trench.

We perfused liquid growth media continuously through the inlet port, which caused most of the population in the main trench to exit from the device. We maintained the chamber at 37°C and recorded time-lapse phase images of individual cells trapped in the orthogonal channels over the next 7 hours. We used an automated microscope stage to repeatedly cycle between different regions of interest within the device. This enabled us to monitor the growth of multiple cells over the duration of our experiment. The recorded images from each region were analyzed to quantify the changes in the lengths of single cells over time (see *Materials and Methods*).

Out of 154 motile cells that we observed, 139 cells grew longer and eventually divided; 15 cells did not grow or divide (**Fig 3A**). Thus, almost 90% of the motile-tolerant cells remained culturable. The single-cell growth data indicated the presence of multiple subgroups within the motile-tolerant population, with some cells resuming growth faster than others.

**Figure 3.**
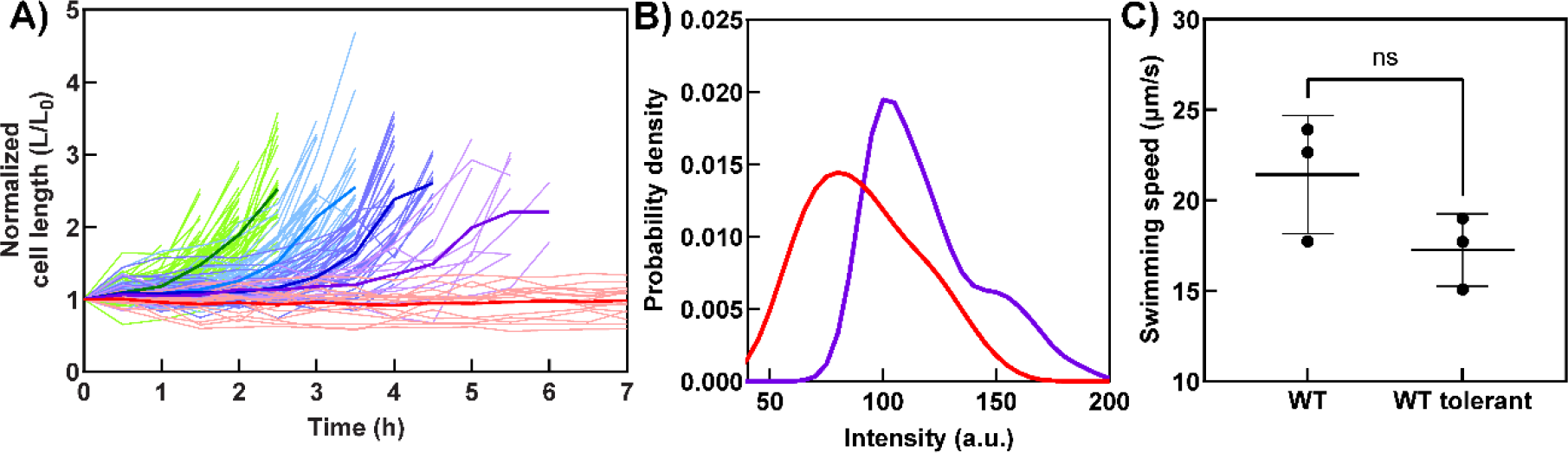
Motile-tolerant cells remain culturable. **A)** Time-varying changes are indicated in cell lengths (L) normalized by their initial lengths (L_0_) at the beginning of observation. Light curves indicate raw values and bold curves indicate mean values calculated for each color group: green curve (n = 62 cells), light blue (n = 37 cells), dark blue (n = 30 cells), and purple curve (n = 10 cells). The red group did not divide during the observation time (n = 15 cells). Approximately 71% of the dividing cells divided within 3.5 h, and the rest exhibited a greater delay in initiating division. **B)** The probability densities are calculated from intensity distributions of the motile-tolerant cells trapped inside the orthogonal channels (blue curve) and the non-motile fraction in the main trench (red curve). The motile-cell mean intensity was 119 ± 24 (n = 151 cells) and the nonmotile-cell mean intensity was 92 ± 24 (n = 84 cells). **C)** Mean swimming speeds of sensitive wild-type was 21.43 ± 3.266 (n = 402 cells) and tolerant wild-type cells was 17.26 ±2.001 (n = 135 cells) determined from three biological replicates (ns indicates P > 0.05 by Welch’s t test).

Thus, a significant majority of the motile-tolerant cells retain the ability to grow. Because motility is a significant fitness factor, these data suggest that motility could play a substantial role in spreading tolerant bacteria.

### Motile-tolerant cells retain high membrane potential

Our finding that most motile-tolerant cells are culturable suggests that they shield themselves from long-term damage by the antibiotic. As this tolerance is not due to the acquisition of resistance genes or mutations (*32*), the survival mechanism could involve some form of dormancy. However, this is improbable, considering motility requires energy reserves that dormant cells are unlikely to possess. A likelier possibility is that the motile-tolerant cells survive by a mechanism powered by the PMF, perhaps by utilizing efflux pumps to decrease the intracellular concentration of the antibiotic (*58*). Alternatively, these cells might survive by limiting the permeation of the antibiotic by actively inhibiting porin activity (**Fig 1**).

To pinpoint the survival mechanism, we compared ΔΨ in both motile and non-motile cells (*33*). We treated the tolerant cells with a membrane-potential reporter dye, thioflavin T (see *Materials and Methods*). This dye labels negatively charged cell membranes, such that when exposed to appropriate irradiation, cells with a greater membrane potential fluoresce brightly (*33*). Next, we isolated the motile fraction in the orthogonal channels of the microfluidic chamber as before. We illuminated the cells and recorded fluorescence emissions with time-lapse imaging (see *Materials and Methods*).

We quantified the fluorescence intensities of the motile fraction by analyzing cells exclusively found in the orthogonal channels. For the non-motile fraction, we quantified intensities by focusing on cells found in the main trench. In this latter analysis, we excluded any motile cells by revisiting the recorded time-lapse fluorescence images and selecting only the cells that appeared to diffuse passively within the field of view. The probability density functions calculated from the intensity distributions of both fractions can be seen in **Fig. 3B** (see raw data in **Fig. S3**). The motile fraction exhibited a higher intensity on average compared to the non-motile fraction. This suggested that the motile-tolerant cells indeed retained a higher membrane potential compared to the non-motile cells. The motile-tolerant cells swim at speeds similar to those of sensitive wild-type cells (**Fig 3C**) (*32*). As the rotational rates of the flagella are proportional to the PMF, this indicates that the motile-tolerant cells have a similar PMF as sensitive wild-type cells.

### Motile-tolerant cells do not rely on efflux alone to survive

Given that the motile-tolerant cells possessed higher membrane potential than the nonmotile cells, we investigated whether the former subgroup survived in the presence of ampicillin by augmenting efflux activity after potentiation with rifampicin. For this purpose, we utilized ethidium bromide (EtBr)– a DNA-intercalating fluorescent dye. EtBr is commonly used to monitor bacterial efflux activity as it is secreted via efflux pumps. A decrease in the efflux activity due to the dissipation of the membrane potential can be readily estimated by monitoring changes in the intracellular fluorescence (*59–63*).

To characterize EtBr’s ability to report efflux and porin functions, we treated sensitive cells of *E. coli* with 1μg/mL EtBr and observed no fluorescence signals (**Fig 4A**). Next, we treated the sensitive cells with 100 μM of an efflux pump inhibitor, PaβN (Phe-Arg β-naphthylamide dihydrochloride), for 30 minutes before resuspending in EtBr (see *Materials and Methods*). PaβN putatively binds to AcrB of AcrAB-TolC efflux pump (*64*), inhibiting efflux (*65*). These complexes form the major efflux system in *E. coli*, conferring protection against ampicillin. We recorded strong signals from the cells. The cells also retained motility. We also dissipated the PMF with an uncoupler, carbonyl cyanide 3-chlorophenylhydrazone (CCCP), again registering strong EtBr signals (**Fig S4**). CCCP-treated cells were non-motile.

**Figure 4.**
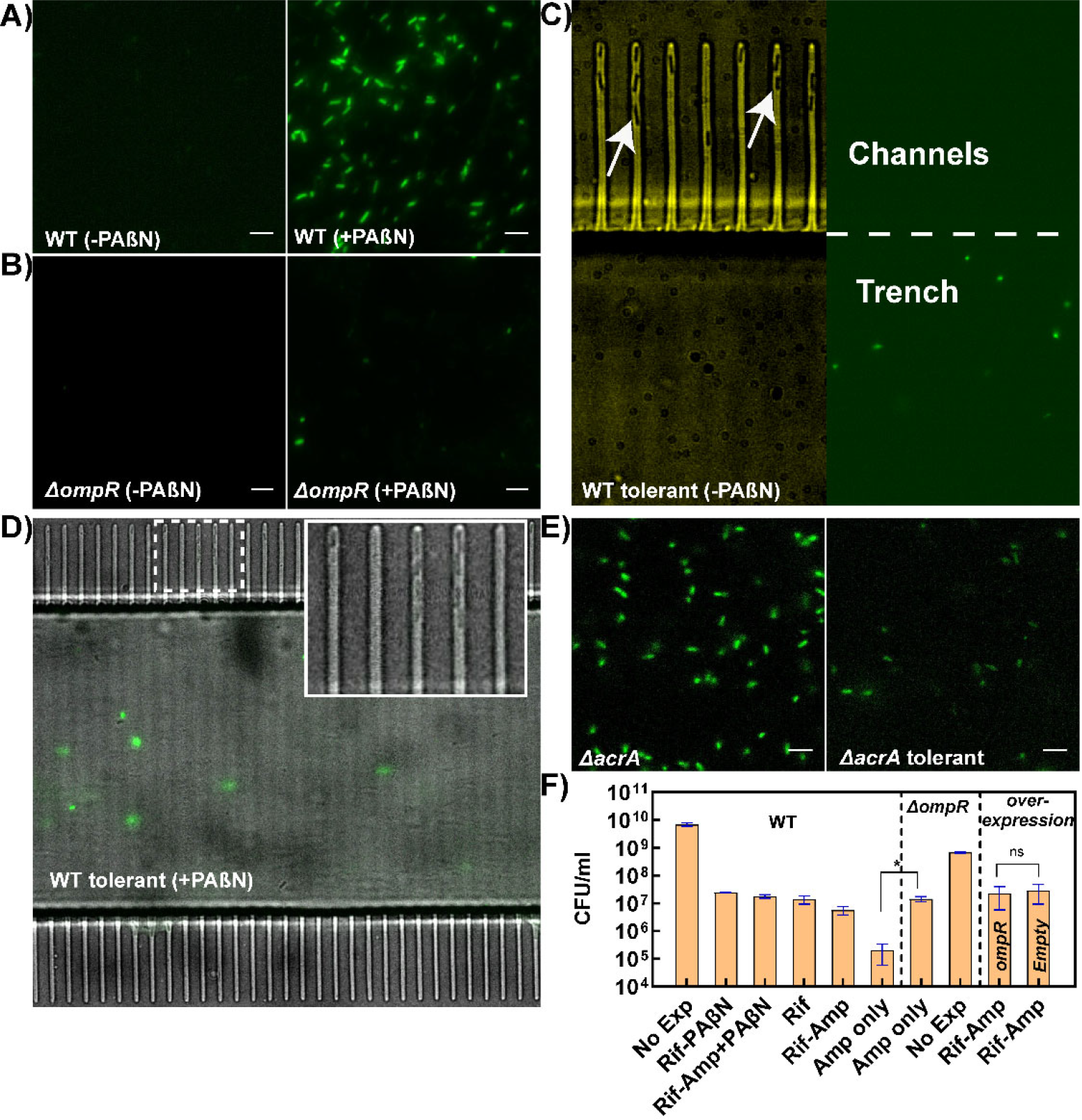
EtBr as a probe for transport. **A)** *Left panel*: Fluorescence data indicate that wild-type cells do not accumulate EtBr intracellularly. *Right panel*. Treatment with PaβN induces EtBr accumulation inside the wild-type cells. **B)** *Left* and *right panels*: OmpF cells mostly fail to accumulate EtBr, with or without PaβN. **C)** *Left panel*: Phase image indicates the presence of several tolerant-motile WT cells in the channels (indicated by the white arrows). *Right panel*: The corresponding fluorescence data revealed signals only from nonmotile cells. The white dotted line indicates the boundary between the orthogonal channels and the trench. **D)** The composite image overlays the phase and fluorescent (green) data for tolerant WT cells treated with PaβN and EtBr. Only the non-motile tolerant cells fluoresced, indicated by the green signal in the main trench. The inset shows the presence of motile-tolerant cells in channels. **E)** *Left panel:* The strong fluorescence signals indicate a significant accumulation of EtBr in the *ΔacrA* cells. *Right panel:* EtBr accumulation was negligible in *ΔacrA* tolerant cells generated with the Rif-Amp treatment. **F)** CFU assays for Wild-type: No-exposure control (6.95 × 10^9^ ± 9.69 × 10^8^ colonies), Rif-PaβN, 3h exposure (2.47 × 10^7^ ± 8.84 × 10^5^colonies), Rif-Amp-PaβN exposure, 3h (1.79 × 10^7^ ± 2.66 × 10^6^ colonies) rifampicin-exposure for 30 min (1.4 × 10^7^ ± 4.75 × 10^6^ colonies), Rif-Amp exposure, 3h (5.6 × 10^6^ ± 1.85 × 10^6^ colonies), and Amp only 3h exposure (2.00 × 10^5^ ± 1.41 × 10^5^ colonies); CFU assays for *ΔompR*: Amp only 3h exposure (1.43 × 10^7^ ± 3.06 × 10^6^ colonies), and No-exposure control (6.92 × 10^8^ ± 4.60 × 10^7^ colonies); CFU assays for overexpression studies with 25μM IPTG: Rif-amp exposure of OmpF overexpression (2.26 × 10^7^ ± 1.68 × 10^7^ colonies) and Empty vector control (2.87 × 10^7^ ± 1.94 × 10^7^ colonies). The mean CFU/mL values were obtained from at least two biological and multiple technical replicates (* indicates P ≤ 0.05 and ns indicates P > 0.05 by Welch’s t test).

Next, we imaged the uptake of EtBr in a mutant lacking the *ompR* gene. OmpR is a response regulator that modulates porin expression, and in its absence, the number of major porins, including OmpF, is significantly decreased (*66*). In this mutant, no EtBr uptake was observed with or without PaβN treatment, indicating that OmpF is necessary for EtBr uptake (**Fig 4B**). Through phase microscopy, we confirmed that the cell density across all these datasets was similar. Therefore, instances where fluorescence data indicated few cells or no signal were attributable to the absence of EtBr accumulation rather than a scarcity of cells within the field of view. Thus, EtBr is an excellent probe to study efflux and porin-mediated transport across the cell membrane (*67, 68*).

We then treated the Rif-Amp-induced tolerant cells of the wild-type strain with EtBr and loaded them into the microfluidic device. The motile-tolerant cells populated the orthogonal channels as before. When illuminated, strong emission signals were observed only in the main trench (**Fig 4C**); none of the motile cells in the orthogonal channels exhibited any fluorescence. By visually inspecting cell movements in the time-lapse data, we confirmed that the fluorescence signals in the main trench were exclusively from non-motile cells. Thus, the motile cells did not appear to accumulate EtBr intracellularly.

To determine if the lack of EtBr accumulation in the motile-tolerant cells was due to increased efflux activity, we performed the PaβN treatment on the tolerant cells. We treated cells potentiated with Rifampicin with 100 μM of PaβN and 100 μg/mL Ampicillin for 2.25 hours (see *Materials and Methods*). We then added EtBr and incubated the cell suspension for an additional 45 minutes before loading the cells into the microfluidic chamber. The motile cells populated the orthogonal channels but did not exhibit any fluorescence signal. Fluorescence emissions were only visible in the trench from nonmotile cells (**Fig 4D**). Considering PaβN successfully inhibits efflux (**Fig 4A**), the lack of EtBr accumulation in the PaβN-treated motile-tolerant cells suggested that their permeability had decreased.

To confirm these results, we worked with an *acrA* mutant strain that lacked the major efflux system. We imaged the uptake of EtBr before and after Rif-Amp treatment. Prior to the Rif-Amp treatment, the *ΔacrA* cells exhibited strong fluorescence, indicating that any residual efflux activity was insufficient to remove the dye from the cytosol. In comparison, Rif-Amp-treated tolerant cells of the *ΔacrA* strain fluoresced weakly when exposed to EtBr (**Fig 4E**). We quantified the difference in fluorescence emission density in the two treatments by calculating the mean intensity per pixel in individual cells (**Fig S5**). The difference in the mean pixel intensity in the tolerant and sensitive *ΔacrA* cells was significant (p-value < 0.001). **Movies 2** and **3** qualitatively illustrate these intensity differences and demonstrate the ability of the tolerant *ΔacrA* cells to swim. Thus, the uptake of EtBr was inhibited in tolerant cells of the *ΔacrA* strain despite the lack of efflux activity, again suggesting decreased permeability in tolerant cells.

Considering EtBr and ampicillin have different molecular properties, it is possible that OmpF permeability remained unchanged for ampicillin even if it decreased for EtBr. If so, we hypothesized that inhibiting efflux activity could decrease tolerance to ampicillin by promoting ampicillin accumulation in the cell. To test this, we exposed Rif-treated WT cells to 100 μM of PaβN and 100 μg/mL Ampicillin for three hours and recorded their motility and viability (CFU). The results were inconsistent with our hypothesis: the cells did not lose motility despite treatment with PaβN (**Movie 4**), and PaβN treatment could not decrease the number of colony-forming units in the tolerant cells (**Fig 4E**).

To confirm that the dissipation of efflux did not mitigate tolerance, we tested the effect of the Rif-Amp treatment on *ΔacrA* cells. These cells swam robustly before the Rif-Amp treatment, but only a fraction remained motile in the tolerant population (**Movie 5**). CFU assays showed that the survival rate of the tolerant *ΔacrA* cells was not lower than the tolerant cells of the wild-type strain. We also attempted to delete *tolC*. However, the strain was poorly motile after Rif treatment (**Movie 6**). Therefore, we did not include this strain in further experimentation.

These results imply that in tolerant cells, the mass transport barriers to Ampicillin are similar to those observed for EtBr (**Fig 4D**). Also, efflux is not necessary for the survival of motile-tolerant cells.

### Motile-tolerant cells inhibit permeability

If decreased permeability is responsible for the lack of EtBr accumulation in the tolerant *ΔacrA* cells, we hypothesized that forcibly perforating the cell via electroporation would eliminate transport barriers to the uptake of EtBr. We electroporated electrocompetent cells of the tolerant *ΔacrA* strain in the presence of EtBr (see Methods). As a control, we electroporated electrocompetent cells of the antibiotic-sensitive wild-type strain. As shown in **Fig 5A**, the sensitive cells of the control population and the tolerant cells of *ΔacrA* fluoresced brightly immediately post-electroporation. The fluorescence signal in the control cells decreased over the next hour as their efflux systems recovered to pump out the EtBr. In comparison, the fluorescence signal did not decrease in the tolerant *ΔacrA* cells in this duration. Thus, EtBr enters the *ΔacrA* cells once mass transfer barriers are overcome and accumulate permanently as the cells lack strong efflux activity.

**Figure 5.**
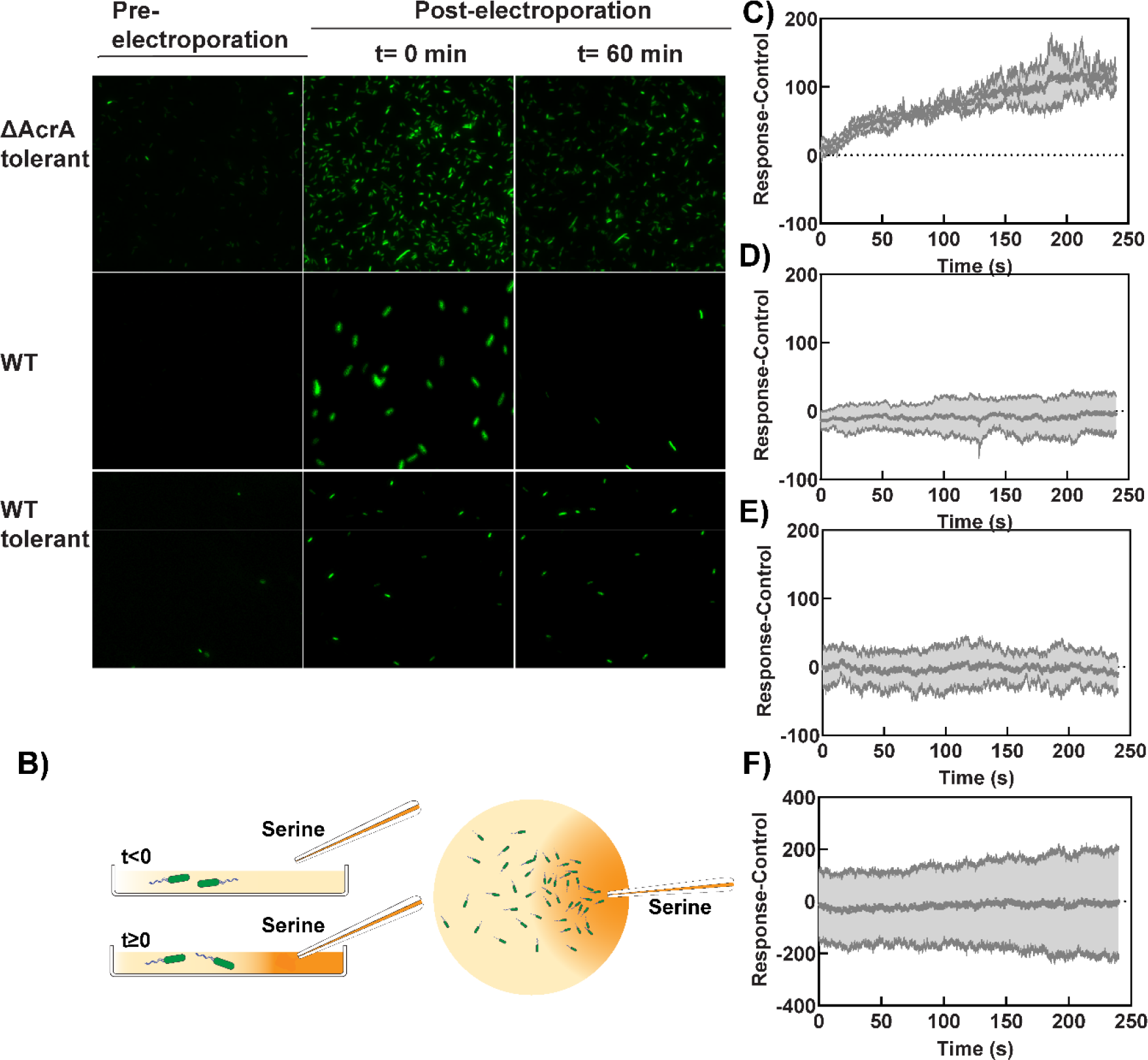
Chemotactic assays reveal inhibition of porin activity. **A)** Comparison of EtBr accumulation in cells permeabilized by electroporation. *Top panel* indicates wild-type cells, the middle panel indicates Rif-Amp treated tolerant wild-type cells, and the *bottom panel* indicates tolerant *ΔacrA* cells. All the cells fluoresced immediately post-electroporation. The signals in the sensitive wild-type cells decreased substantially over an hour but not in the tolerant wild-type or *ΔacrA* cells. **B)** Schematic shows a reservoir containing motile *E. coli*. A micropipette containing L-serine (1mM) is lowered into the reservoir to apply a chemical stimulus at *t = 0* s. The chemotaxis response is measured from the variation in cell density near the micropipette tip over time. **C** to **F)** The density of normal cells (sensitive cells) increased rapidly near the micropipette tip when it contained serine (Response) but not when it contained MB (Control). The ordinate indicates the difference between the cell densities during the Response and Control measurements. The mean response was calculated over three biological replicates. **C)** Wild-type cells exhibited a strong chemotaxis response to serine. In contrast, an attenuated chemotaxis response was observed in **D)** tolerant wild-type cells (Rif-treated cells exposed to Amp for 3 h), **E)** tolerant wild-type cells (Rif-treated but not exposed to Amp), and **F)** OmpR-deleted strains with no exposure to either Rifampicin or Ampicillin.

We repeated the electroporation experiments with Rif-Amp-treated tolerant cells of the wild-type strain. The cell density was lower in the field of view as the tolerant fraction is relatively small. Nevertheless, fluorescence imaging revealed negligible accumulation of EtBr prior to electroporation, as anticipated (**Fig 5A**). Post-electroporation, we measured strong signals in cells lasting for almost an hour. This was consistent with our hypothesis that the tolerant cells inhibit porin function to limit transport across the outer membrane.

If Rif-treated tolerant cells inhibit porins to prevent Ampicillin accumulation, what would occur if Ampicillin entered the cells prior to Rif potentiation? To investigate, we exposed wild-type cells to Ampicillin for 5 minutes and subsequently administered the standard Rif-Amp treatment for 3 hours. CFU counts decreased by an order of magnitude as Ampicillin entered the cells before they could inhibit porin activity, in line with our expectations.

### Inhibited porin function promotes survival in tolerant cells

If the tolerant cells indeed survive by modulating their permeability to antibiotics, the mechanism likely involves inhibition of general porins such as OmpF, as they mediate the transport of antibiotics into the cell (*69–71*). In that scenario, a strain lacking porins would be able to tolerate the 3 h Amp treatment without needing potentiation with rifampicin. To test this, we used the strain deleted for *ompR*.

We directly exposed the *ΔompR* cells to ampicillin for 3 h, skipping the Rif treatment. As anticipated, the porin-deficient mutant exhibited a higher survival rate than the control wild-type strain that was exposed to ampicillin for 3 h directly (**Fig 4F**). The survival rates were somewhat higher than those of the wild-type strain’s tolerant cells (**Fig 4F**), consistent with the idea that the loss of porin function is likely the cause of tolerance adaptation in wild-type cells.

To determine if the mechanism of porin inhibition involved the downregulation of porin-related genes, we overexpressed the OmpF from a high-copy number, inducible plasmid in the wild-type strain (see *Materials and Methods*). The rationale was that increasing the expression of porins might compensate for decreased synthesis during the 30-minute Rifampicin exposure, rendering transcriptional or post-transcriptional adaptation mechanisms less effective. We repeated the Rif-Amp treatment with the overexpression strain and compared the survival against a control wild-type strain that carried the empty vector. Increased porin expression did not decrease the survival of the tolerant population **(Fig 4F)**. Although several tolerant cells with overexpressed OmpF were observed to tether and rotate, indicating that they retained flagellar function, there were very few swimmers (**Movie 7**).

Thus, the overexpression of porins did not decrease the tolerance induced by Rif-Amp treatment. Based on our results, it is likely that the porin adaptation mechanism is independent of alterations in gene expression. Rather, the mechanism probably alters the permeability of existing porins in the outer membrane, for example, by altering charged residues within each pore by modulating periplasmic pH. If so, it would explain why increased expression of porins does not impact tolerance.

### Porin adaptation prevents small molecule transport

Measuring the decreased transport of ampicillin in tolerant cells is challenging. For example, we attempted to measure the transport of Bocillin, a fluorescent derivative of ampicillin. However, its size is greater than 600 Da, which is the cutoff for OmpF porins. Hence, few or no cells took up the compound even when efflux was dissipated. Similarly, it is challenging to measure ampicillin uptake with mass spectrometry (LC-MS/MS) as it covalently binds to its target (*72, 73*).

We attempted to circumvent these technical limitations by measuring the transport of a molecular smaller than ampicillin, L-serine. Serine diffuses into the periplasm via the porins and interacts with the periplasmic domains of the chemoreceptors situated in the inner membrane (**Fig. 1**). This induces a strong positive chemotaxis response through the modulation of flagellar rotation, causing cells to swim toward higher serine concentrations (*74*). We reasoned that if the porins are constricted or blocked in the motile-tolerant cells, serine would fail to reach the receptors, leading to an attenuated chemotaxis response.

Guided by the aforementioned rationale, we utilized a chemotaxis assay to compare the differences in porin activity in normal and tolerant cells. We employed a 6 μm diameter micro-capillary, controlled by an automated micromanipulator, to administer a small dose of 1 mM serine in a petridish filled with buffer containing swimming cells (**Fig 5B**). Cell migration was recorded using phase microscopy, and the response to serine or the control (buffer) was quantified using cell-tracking algorithms (*32*). We applied speed-based filters to eliminate non-motile-tolerant cells observed in the region of interest from our analysis (*Materials and Methods*).

When stimulating antibiotic-sensitive wild-type cells with serine, we observed a significant increase in the motile cell density near the micro-capillary tip within a few minutes of the stimulus. The cells did not respond when buffer solutions were introduced via the micro-capillary in control experiments. We quantified the net chemotaxis response by subtracting the control densities from the response densities over the duration of observation (**Fig 5C**). As evident, there was a rapid increase in cell densities near the source of serine.

In comparison, *motile-tolerant cells* that were generated with the Rif-Amp treatment did not exhibit a significant chemotaxis response to serine (**Fig 5D**). It is noteworthy that while the tolerant-motile cells retained motility, they were unable to undergo chemotaxis. To determine whether the lack of chemotaxis response was due to changes in receptor expression levels induced during the 3 h exposure to ampicillin, we performed the assay with motile cells treated for 30 minutes with only rifampicin. We observed a similar attenuated response in Rif-treated motile cells (**Fig 5E**), indicating that rifampicin exposure inhibited the porins. As a final control, we measured the chemotaxis response in the motile fraction of the OmpR mutant cells (**Fig 5F**). Again, no chemotaxis was observed. These observations are consistent with the notion that serine transport through the porins is greatly diminished in tolerant cells.

Overall, our results suggest that the 30-minute rifampicin exposure significantly attenuates porin-mediated ligand transport in the motile-tolerant cells.

### Transport model predicts that motile-tolerant cells survive by porin inhibition and non-motile tolerant cells survive via dormancy

Although the motile-tolerant cells retain a high PMF like the sensitive population (**Fig 3B**), efflux alone is inadequate in preventing drug accumulation and cell death. Evidently, rifampicin induces a rapid decrease in porin function within a few minutes. How effective is the porin inhibition mechanism in preventing drug accumulation, and what is the relative importance of efflux versus porin-mediated transport in antibiotic tolerance? To address these questions, we employed the following antibiotic transport model.

#### Model and experimental determination of kinetic parameters

For a cell with volume V, the transport of a ligand across the cell boundary can be represented by the following equation.

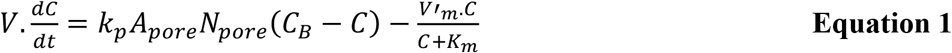

Here, the first term on the right-hand side represents passive diffusion of the ligand into the cell due to a concentration gradient across the cell boundary, and the second term represents the PMF-powered efflux of that ligand to the extracellular environment. *C* represents the intracellular concentration of free ligand at an arbitrary time, *t*. We reasonably assume that the drug is not degraded inside the cell. We maintained a bulk concentration *C*_*B*_ = 1 μg/mL in all our experiments.

Briefly, our approach involved experimentally determining the basal values of parameters controlling efflux and passive influx, separately, in antibiotic-sensitive wild-type cells. Then, we determined how these parameter values deviate from their basal values in the motile-tolerant and non-motile tolerant subtypes to determine their survival mechanisms.

To determine the parameters controlling passive influx, we eliminated efflux in wild-type cells with PaβN treatment for an hour such that *V′*_*m*_ *=* 0 in equation 1. These cells possessed normal porin function, and an hour-long treatment was adequate to equilibrate the ligand concentration, 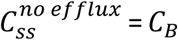 (= 1 μg/mL). We illuminated cells trapped between a soft-agar pad and the coverslip and recorded their emissions. From the fluorescence images, we calculated the single-cell mean intensity 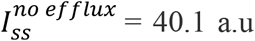 (see **Fig 6A**), corresponding to 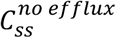. This yielded the conversion factor, *I*_*c*_ = 40.1 a.u.(mL/μg), which related the fluorescence intensity per cell to the intracellular EtBr concentration:

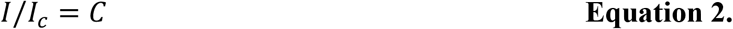

**Figure 6.**
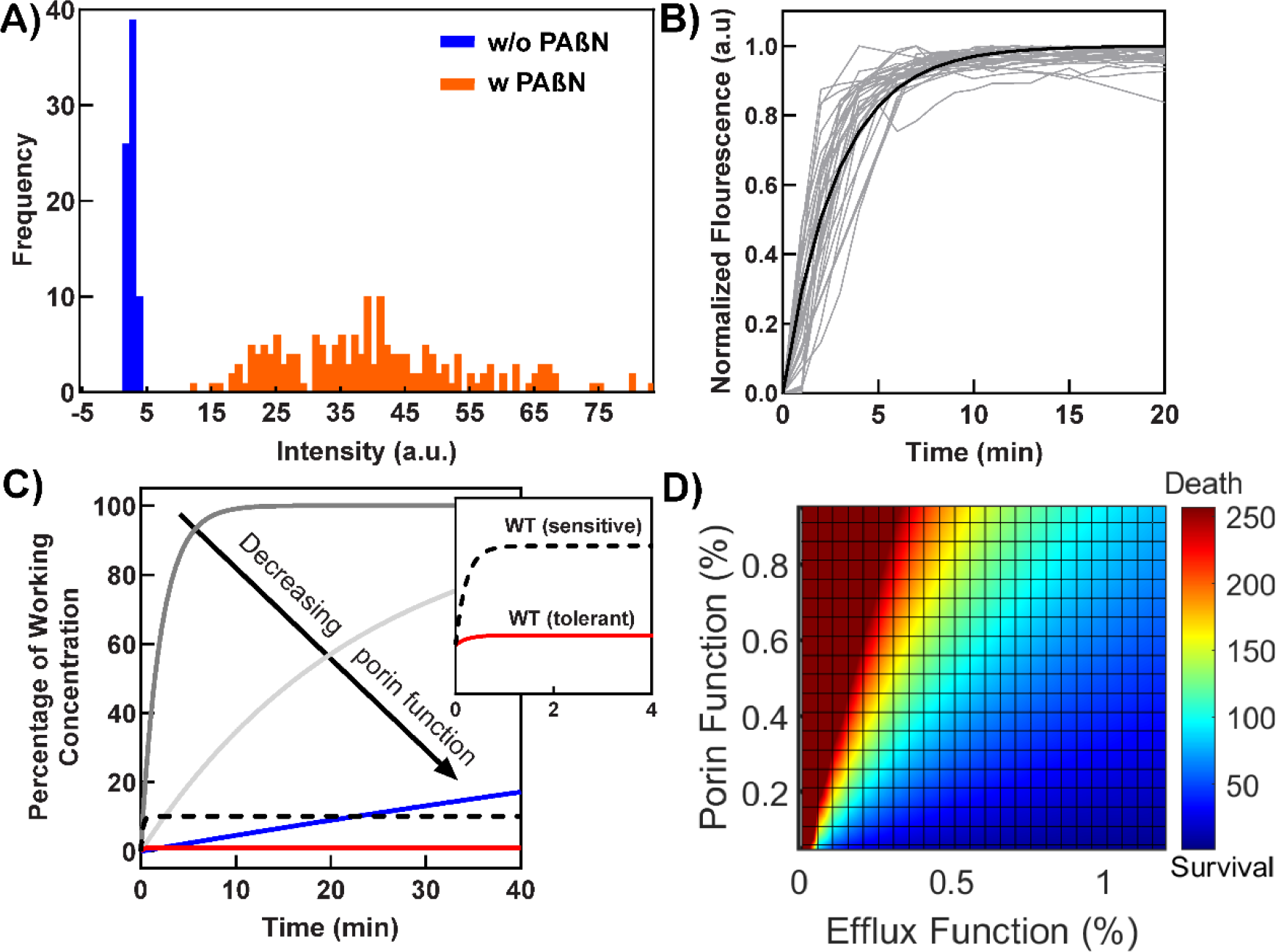
EtBr uptake kinetics. **A)** The distributions indicate the pixel intensities in individual cells of the wild-type strain in the presence of EtBr. The fluorescence emissions were higher in cells treated with PaβN (orange data, n = 175, mean pixel intensity = 40.1 +/-14.6 a.u.) compared to cells that were not exposed to PaβN (blue data, n = 75, mean pixel intensity = 2.8+/-0.6 a.u.). **B)** The gray emission curves indicate EtBr uptake in individual cells of the wild-type cells treated with PaβN (n = 30 cells). The individual curves were scaled and offset to originate at *t* = 0 min (gray curves). The fit equation, *Î* =1-exp(-*k*_1_.*t*) and the scaled data were linearized and the mean value of *k*_1_= 0.47 min^-1^ (0.43, 0.5 min^-1^, 95% confidence interval) was calculated from linear regression. The black curve shows the fit. The raw data are shown in Fig S5. **C)** Curves show predicted kinetics of drug uptake in individual cells from equation 1. The black dotted curve represents uptake in wild-type cells that are sensitive to the drug. The motile-tolerant cells (red curve) retain much lower intracellular concentrations due to decreased porin activity: 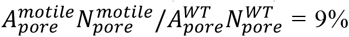. Inset shows the time for equilibration in the two populations. Nonmotile-tolerant cells lacking efflux activity can adapt by decreasing porin activity to limit the rate of drug uptake: gray curve = 0%, light gray = 90%, and blue curve = 99% decrease in porin function. **D)** The steady-state intracellular drug accumulation due to antibiotic adaptation is shown relative to the baseline steady-state in cells that cannot adapt. A value of 100 indicates wild-type cells that perish under antibiotic stress. Tolerant cells adapt via porin inhibition to survive (dark blue zones), provided they can support residual efflux function with PMF. Efflux function > 1 indicates hyper-polarized cells.

Next, we estimated the rate at which the intracellular concentration in individual cells equilibrated during the hour-long PaβN treatment. As shown in **Fig S6**, individual cells lost their efflux activity at different times, indicated by a sudden and rapid increase in intensity, i.e., EtBr uptake. Solving **Eq. 1** with *V′*_*m*_ *=* 0 and combining it with **Eq. 2** yielded 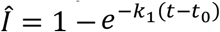, where 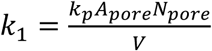 and *Î* is the scaled intensity. Fits to experimental intensity curves (**Fig 6B** and **Fig S7**) yielded *k*_1_= 0.47 min^-1^. We reasonably assume that the permeation through the outer and inner membranes can be represented by a single parameter, *k*_*p*_. Based on literature values, *k*_*p*_=3×10^−6^ m/s (*75*), the diameter of OmpF porin pore =0.7 nm (*76*), and, cell length =2.5 μm and width = 0.5 μm (*51*). From the fitted value, we estimate the number of pores, *N*_*pore*_∼ 1.7×10^4^ in our cells. This correctly falls within the range reported for porins (*77*).

Having determined the influx parameters, we quantified the ratio of the two parameters for efflux, 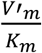, in wild-type cells possessing normal efflux activity and porin function. To do this, we measured the mean fluorescence intensity in wild-type cells treated with only EtBr for an hour. At steady state, equation 1 combined with **Eq. 2** yields:

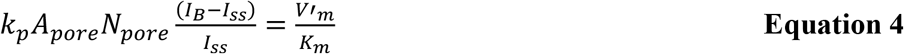

Here, we reasonably approximated that the intracellular buildup of an antibiotic is much less than *K*_*m*_, i.e., *C*_*ss*_ ≪ *K*_*m*_, based on previous work (*75*). Our analysis of fluorescence images indicated *I*_*ss*_ = 2.8 a.u (**Fig 6A**). From **Eq. 4**, we obtained 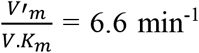. For a nominal value of *K*_*m*_= 200 μM for antibiotics (*75*), *V′*_*m*_/*V*= 1.3 mM/min, where *V* is the cell volume.

Finally, we measured the mean cell intensity in wild-type cells made tolerant by treatment with Rifampicin and suspended in EtBr for 1 hour. As the emissions from the tolerant cells were weak, we employed a highly sensitive EMCCD camera to boost the signal-to-noise ratio. The fluorescence signals from the motile-tolerant cells were indistinguishable from cell auto-fluorescence, which is about ∼10% of the emissions from sensitive wild-type cells. This provided an upper bound for EtBr uptake in the motile-tolerant cells. Plugging this value in **Eq. 4**, and making a reasonable assumption that the efflux activity in the two populations is similar, we calculated the upper bound for the overall porin area in the motile-tolerant population: 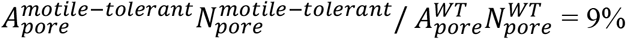 or less.

Our OmpF-overexpression measurements suggest that the tolerance mechanism is independent of the number of OmpF porins. Hence, we assume that there is no decrease in the number of porins in the tolerant cells relevant to the sensitive cells, i.e., 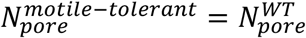. In that case, our calculations indicate that individual porins in the motile cells constrict during the adaptation to antibiotic stress, i.e., 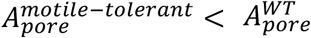. This occurs due to a decrease in the effective diameter of each porin from ∼ 0.7 nm to 0.2 nm or less in the motile-tolerant cells. While the physical diameter of the pore might remain unchanged, the effective diameter probably decreases due to a modified electrostatic charge distribution. These modifications likely repel charged extracellular species like ampicillin.

In comparison, we detected strong emissions from the non-motile tolerant cells, indicating they lack sufficient efflux to expel EtBr (**Fig 4C, D**). This observation is consistent with the observed decrease in membrane potential within these nonmotile cells (**Fig 3B**). Thus, these cells are probably unable to prevent antibiotic accumulation and are likely to rely on dormancy as a strategy for survival.

#### Model predictions

From the measured kinetic parameters, we predicted the rate of intracellular drug accumulation in cells that are exposed to an antibiotic at time t = 0 s. In sensitive cells and in motile-tolerant cells, the intracellular antibiotic levels reach a steady state within a few seconds (**Fig 6C**). The rapid equilibration is dominated by efflux kinetics (6.6 min^-1^) as the efflux rate is almost an order of magnitude higher than the rate of passive influx (0.47 min^-1^). Significantly, the steady-state intracellular concentration in sensitive cells is above the level needed to kill them. Because of decreased porin function, the corresponding concentration in motile-tolerant cells is an order of magnitude lower, allowing them to survive.

The nonmotile-tolerant cells have lower membrane potential than the motile-tolerant cells, and, therefore, have depressed efflux activity. Hence, we explored the consequences of the loss of PMF and the associated efflux function in these cells. As shown in **Fig 6C**, the intracellular drug concentration in cells with no efflux activity required several minutes to equilibrate if they had normal porin function (red curve). The time needed for equilibration continues to increase in such cells with decreasing porin function **(Fig 6C**). However, eventually, the drug will accumulate in these cells, and therefore, they probably need to survive by inducing cell dormancy.

While slow kinetics of accumulation certainly improve the chances of survival by enabling extra time for adaptation, the steady-state concentration of the accumulated drug strongly influences cell fate. If the steady-state intracellular concentration of the drug is higher than the threshold concentration, the chances of survival are low in non-dormant cells. Solving equation 1 at steady state 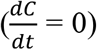, we calculated the dependence of steady-state intracellular drug concentration on the relative rates of efflux-based drug expulsion and passive influx via the porins.

In **Fig 6D**, we show the intracellular drug accumulation at a steady state in cells relative to the accumulated amount in wild-type cells with normal efflux and porin function. A value of 100 or above (light blue zone) indicates cell death. Wild-type cells do not survive without Rifampicin-potentiation because ampicillin concentration exceeds tolerable levels, despite possessing normal efflux and porin function. As discovered in this work, some cells can survive with normal efflux function provided they can adaptively inhibit porin function upon Rifampicin exposure (blue zones). Hyperpolarization coupled with porin inhibition boosts survival chances significantly by substantially decreasing drug accumulation. Finally, a complete loss of PMF results in cell death (orange/red zones), unless the cells achieve dormancy by inducing stress-response mechanisms.

The adaptive mechanism of porin constriction is likely to involve charge redistribution or alterations in the electrostatic environment within each porin, thereby decreasing the permeability of different ligands such as ampicillin, serine, and EtBr. Such a mechanism is consistent with the observation that overexpression of OmpF does not inhibit tolerance.

## Discussion

It is common for bacteria stimulated with one antibiotic to become tolerant to a different one (*34, 53*). Rifampicin inhibits transcription and decreases cellular respiration rates (*34*), reportedly inducing cells to become antibiotic-tolerant through dormancy. Although dormant cells lacking in PMF can neither power efflux nor divide to dilute existing porins, our discovery of a sub-population of tolerant cells with high PMF suggested that these cells could survive by decreasing the intracellular buildup of antibiotics with efflux pumps (*32*). Owing to significant phenotypic heterogeneity in tolerant populations, it was necessary to segregate these high PMF motile cells to test the efflux hypothesis. Our serendipitous finding that motile cells penetrate confined spaces exclusively enabled us to segregate them from a mixture of motile and non-motile tolerant populations and probe their physiology at a single-cell level.

We observed that ∼ 90% of the motile-tolerant population was viable and culturable, as evidenced by their ability to grow and divide once the antibiotic stress diminished. The resuscitation or ‘awakening’ rate following the antibiotic removal varied significantly within the motile population, consistent with previous reports of high heterogeneity in the rate at which cellular growth is recovered by persister-like cells (*78*). Motility is a major fitness factor, and therefore, motile-tolerant cells are biomedically significant as they can rapidly spread to antibiotic-free areas.

Cells can prevent the accumulation of antibiotics by expelling them with PMF-powered efflux pumps faster than the rate at which they enter through passive diffusion through the porins. We hypothesized that if the high-PMF motile-tolerant cells relied on efflux pumps for survival, inhibiting efflux would selectively eliminate the motile cells in the presence of ampicillin. We tested this notion by inhibiting efflux, either with PaβN or by deleting *acrA*, a major component of efflux pumps in *E. coli*. We observed no decline in cell survival or motility in tolerant populations (**Fig 4F**), consistent with the idea that ampicillin intake itself was decreased in tolerant cells.

We performed single-cell fluorescence microscopy experiments on motile-tolerant cells to quantify intracellular accumulation of Ethidium bromide (EtBr), which we showed was capable of reporting efflux activity and porin function (**Fig 4A, B**). The motile-tolerant cells did not accumulate EtBr. Motile-tolerant cells deficient in efflux activity, either through PaβN treatment or *acrA* deletion, also prevented EtBr accumulation (**Fig 4C, D**). Only forcible perforation with strong electric fields could induce EtBr accumulation in tolerant cells. This was consistent with the idea that tolerant cells inhibited porin function to limit the accumulation of drugs, rather than elevating efflux function. Unlike the motile-tolerant cells, non-motile-tolerant cells accumulated higher levels of intracellular EtBr, presumably relying on dormancy-based survival mechanisms.

A limitation of our studies with EtBr is that it has a similar molecular weight as Ampicillin but distinct chemical properties. These differences could cause variations in porin permeability for each compound in tolerant cells, potentially limiting the extrapolation of our EtBr results to understand Ampicillin transport. These concerns are alleviated by the CFU results in **Fig 4F**, suggesting that Ampicillin intake is indeed restricted in tolerant cells. Nonetheless, we performed an alternate test of our hypothesis by measuring the transport of L-serine, a powerful chemo-attractant that readily enters through the outer membrane porin, in motile-tolerant cells. Serine is smaller than Ampicillin and is sensed by the chemoreceptors situated in the inner membrane (*79*). We hypothesized that if the porins were indeed constricted in the motile-tolerant cells, serine would fail to penetrate the outer membrane and interact with the chemoreceptors. Our results were consistent with this notion; the motile-tolerant cells failed to chemotax toward serine after only 30 minutes of exposure to rifampicin whereas antibiotic-sensitive cells exhibited a strong chemotactic response. A control OmpR strain lacking major porins also failed to respond to serine chemotactically (**Fig 5C-F**).

Although there has been much focus on gene regulatory pathways in antibiotic tolerance, such approaches may fail to reveal short-term adaptations that often underpin initial responses to environmental stressors (*80*). Inhibiting porin-mediated uptake is the first line of defense against antibiotics in Gram-negative bacteria. A delay in antibiotic buildup or a decrease in the steady-state antibiotic levels can provide adequate time for the cell to survive and induce stress-response pathways. Therefore, understanding the kinetics of porin adaptation is paramount.

We mathematically modeled the competing processes of porin and efflux-mediated transport and determined their relative importance in antibiotic tolerance. We measured the kinetic parameters for the two processes experimentally, finding that efflux rates were an order of magnitude faster than porin-mediated intake, and dominated transport in wild-type sensitive cells. Nevertheless, efflux function fails to prevent the intracellular buildup of lethal antibiotic concentrations – without potentiation with rifampicin, wild-type cells succumb in the presence of ampicillin.

Our model indicates that following Rifampicin exposure, a small fraction of the original wild-type population likely constricts OmpF porin diameters from 0.7 nm to ∼0.2 nm within minutes. These motile-tolerant cells survive subsequent antibiotic stress as they couple normal PMF-powered efflux function with reduced porin function to limit intracellular drug accumulation significantly (**Fig 6D**). Another subpopulation likely becomes dormant during Rifampicin exposure and loses motility owing to the loss of respiration. Some of these non-motile tolerant cells probably survive through well-known stress response pathways that do not rely on decreasing intracellular antibiotic build-up (*81–85*).

Our findings suggest that cells employ a nuanced mechanism to decrease the intracellular buildup of antibiotics, balancing porin function to complement efflux in energetic tolerant cells. The porin inhibition mechanism identified in this work is unlikely to depend on the downregulation of porin synthesis, as observed in the OmpF over-expression results (**Fig 4F**), and because Rifampicin is a transcriptional inhibitor.

One possibility is that rifampicin exposure eliminates most of the population except those cells with intrinsically low outer membrane permeability (*86, 87*). But, if this were so, the uptake of nutrients would be inherently low in such cells, to begin with, and motility would be inhibited owing to the lack of resources to synthesize flagellar proteins. However, the motile-tolerant cells are robust swimmers (**Fig 3C)**(*32*). A likelier possibility is that rifampicin-induced stress alters the periplasmic pH, globally constricting extant porins in the outer membrane (*88–90*). Determining the mechanism underlying such short-time porin adaptation is expected to reveal vital insights into antibiotic tolerance in Gram-negative bacterial species.

## Acknowledgments

Microfabrication was performed with the assistance of the Nanoscale Research Facility at the University of Florida, Gainesville. We acknowledge inputs from D. Siegele and Kajol Harsh.

## Data and materials availability

All data are available in the main text or the supplementary materials.

## Materials and Methods

### Bacterial strains, cell culturing and plasmids

Experiments were performed with the *E. coli* RP437 strain (wild-type), or its derivatives, a Δ*cheY* mutant (smooth-swimmer), a Δ*fliC* mutant (nonmotile strain), and *ΔacrA* mutant (efflux-deficient strain). We generated the *ΔacrA* knockout with the aid of the λ-Red mediated homologous recombination technique (*91*). We amplified 2895 bp PCR product from *ΔacrA* Keio knockout collection (Strain JW0452-3) containing kanamycin resistance cassette flanked by FRT sites and additional 300-600bp flanking chromosomal sequence. The PCR product was amplified by following primers: - del_acr_flank_LP 5’-CGTTAAGCTGTTGGCCTTTC-3’ and del_acr_flank_RP 5’-CGCAGTGAACCAGAATAGCA-3’ followed by gel purification. The purified product was transformed in pKD46 containing RP437 strain. After successful genomic integration, the kanamycin cassette was eliminated using pCP20 plasmid. The kanamycin-sensitive strain was sequenced to verify the deletion of *acrA*. We used *ΔompR* Keio knockout collection (BW25113 strain background) directly for the experiments. We retrieved pCA24N-ompF plasmid from ASKA collection (*92*). We excised *ompF* from the plasmid using Sfi-I, gel purified the cut vector and re-ligated, yielding empty pCA24N vector. We used 25 μM IPTG for the overexpression experiments.

We streaked fresh colonies on Luria-Bertani (LB) agar plates from frozen glycerol stocks. We grew overnight cultures in tryptone broth (TB) at 30°C and 40 RPM in a rotating test-tube holder (Fischer Scientific) and diluted the cultures 1:100 times in 10 mL fresh TB the subsequent day to prepare day cultures. We grew the day cultures at 33°C in a shaker incubator set at 170 RPM and harvested the cells once they reached an optical density at 600 nm (OD_600_) between 0.5-0.6, measured with Thermo Spectronic 200 spectrophotometer. The media was supplemented with antibiotics (100 μg/mL ampicillin sodium salt, Sigma Aldrich) where appropriate and fluorescent proteins were expressed by adding 100 μM IPTG. The fluorescent protein genes (eYFP and eCFP) were separately cloned in *ptrc99A* between EcoRI and XbaI cloning sites.

### Generation of tolerant cells

Tolerance was induced by sequentially exposing cells to rifampicin and ampicillin (*32*). Briefly, 5 mL of cells at OD_600_ ∼ 0.8 were collected in a test tube. The cells were treated with rifampicin (Sigma Aldrich) to a final concentration of 100 μg/mL for 30 min, followed by centrifugation. The supernatant was discarded, and the pellet was gently resuspended in 5 mL fresh TB. We added ampicillin to this sample to a final concentration of 100 μg/mL. The cell culture was incubated at 33°C with shaking at 250 RPM for 3 h.

### Motility assays

We washed the cell culture twice in motility buffer (MB) at 1500 g for 5 min in a centrifugation unit (Scilogex SCT412). The supernatant was discarded, and the cell pellet was gently resuspended in an appropriate volume of MB to obtain the desired OD_600_ value. Care was taken during washing and resuspension to prevent the breakage of flagellar filaments. MB was prepared with 0.01 M potassium phosphate, 0.067 M NaCl, 0.1 mM EDTA, 1 μM methionine, and 10 mM lactic acid, and adjusted to pH 7.0.

### EtBr accumulation and membrane potential measurements

We resuspended the tolerant cell pellet in phosphate-buffered saline (PBS) containing 1 μg/mL ethidium bromide (EtBr). The mixture was incubated for 45 min at room temperature, followed by a wash in PBS to remove residual EtBr. The pellet was resuspended in PBS and then loaded into the microfluidic device for imaging. A control vial was prepared in a similar manner, except we supplemented it with carbonyl cyanide m-chlorophenyl hydrazine (CCCP) to a final concentration of 25 μM.

To quantify the kinetics of EtBr uptake, we grew cells to an OD_600_ of 0.5-0.6. We washed and resuspended the cells in TB to concentrate them ∼ 3 folds. We added ∼ 20-30 μl of cells in a 35mm glass bottom microwell dish (MatTek Corp.) and immobilized the cells with the help of 0.8% agarose pad. The agarose pad contained TB, 100 μM PaβN and EtBr(1μg/mL). The cells were imaged subsequently.

We employed electroporation to increase the permeability of the tolerant cells and internalize EtBr. We took 80 μl aliquots of electrocompetent cells and 1ul of 10mg/ml EtBr in a pre-chilled 2-mm electroporation cuvette (Fisherbrand). The electroporation was performed at 1240V in BTX electroporator (Harvard Apparatus). The cells were immediately resuspended in motility buffer containing 1 μg/mL EtBr and imaged.

To measure the membrane potential, we treated cells with thioflavin T (ThT) dye, following previous protocols (*33*). For these experiments, cell pellets were resuspended in 10 mL MB supplemented with 10 μM ThT (MB-ThT solution). We centrifuged the cells again and the pellet was resuspended in MB-ThT to a desired concentration before loading in the microfluidic device.

### Fluorescence and phase imaging

We used a 60x oil immersion TIRF objective on a Nikon Ti-E microscope to obtain simultaneous phase and fluorescence data. The halogen light for phase imaging was filtered with a long-pass filter (720 LP, Chroma) to prevent blue wavelengths from adversely impacting cell physiology. Phase images were recorded with a CCD camera (UI-3240LE, IDS Imaging). Fluorescence images were collected in the epifluorescence mode. Cells expressing eCFP or eYFP were illuminated with a white-LED source (SOLA SE II 365 Light Engine, Lumencor), filtered appropriately: 435/20 ex and 480/40 em (Nikon Inc.) for CFP and ZT514/10, ET520 LP, and ET555/55m (Chroma Inc) for eYFP. Two-channel fluorescence detection was performed in tandem on a sCMOS camera (Andor Zyla, Oxford Instruments) at 2 frames per second with an exposure time of 0.5 s.

For EtBr measurements, we used a combination of 514/10 nm ex filter (Chroma) and a 598/25 nm em filter (Semrock). For ThT measurements, we combined the 435/20 nm ex (Nikon) with a 525/50 nm em (AVR Optics).

### Cell growth and channel occupancy measurements

Time-lapse phase data was analyzed with ImageJ to measure changes in cell lengths. We manually counted empty and occupied channels in the fluorescence data to determine the number of occupied channels and cells per channel.

### Analysis of single-cell fluorescence signals

We fitted elliptical masks on individual cell intensity profiles in ImageJ and totaled the pixel intensities to estimate cell intensities. We confirmed the analysis with custom-written MATLAB codes that subtracted the background intensities from individual cell intensities. For the purposes of illustration in figures, we contrast-enhanced representative fluorescence images, corrected for background, and employed image rotation and false color in ImageJ.

### Microfluidic device operation

The microfluidic device was passivated with 20 mg/mL bovine serum albumin (VWR Life Sciences) for 1 h at room temperature before use. Before each experiment, the air was removed from the device by filling the interiors with MB using a syringe (BD 3 mL) attached to a metal tube (New England small tube corporation 0.28” OD X .016” ID). Approximately 200-300 μL of the washed bacteria were then aspirated into a syringe and connected to the inlet port of the device with a 7-10 cm long polyethylene tubing (0.58 mm inner diameter). The cell concentration for all experiments was adjusted to OD_600_ ∼ 1.3 before loading into the device. The cells were loaded into the main trench from the syringe with a syringe pump (Fusion 200, Chemyx) operated at a flow rate of 200 μL/h. The main trench was visually scanned on a Nikon Ti-E phase microscope with a 20x phase objective. We stopped the flow once the cells were detected in the main trench. To study regrowth, the tolerant cells were loaded using the same method as above. Once the appropriate number of cells entered the channels, we stopped introducing cell suspension and switched to liquid LB at a flow rate of 250 μL/h. The liquid LB was continuously perfused for several hours, and the microfluidic device was maintained at 37°C during the experiment. We repeatedly cycled between channels in different regions using an automated stage (MS-2000, Applied (Scientific Instrumentation) and recorded videos in phase.

### Chemotaxis assay

The tolerant or wild-type cells were diluted in Tryptone Broth (TB) and imaged in a culture dish (Delta T Culture Dish, Bioptechs) with a 20X objective. Motility was recorded for 5 min with a CCD camera (UI-3240LE, IDS Imaging) at 20 frames per second, prior to any stimulus. We introduced a piezo drill tip (Eppendorf) containing either serine or MB (control) into the culture dish with the aid of a micromanipulator (TransferMan 4r, Eppendorf). Post-stimulus motile response was recorded for 5 min at the same frame rate. We analyzed videos with standard particle tracking methods to count the number of cells in the field of view (*48*).

### Statistical analysis

The Student’s t-test was used for all statistical analyses, adjusting for equal or unequal variances as needed. We employed GraphPad Prism, v9.3.0, for analyses and plotting.

